# Epigenetic mediation of AKT1 rs1130233’s effect on delta-9-tetrahydrocannabinol-induced medial temporal function during fear processing

**DOI:** 10.1101/2021.06.30.450484

**Authors:** Grace Blest-Hopley, Marco Colizzi, Diana Prata, Vincent Giampietro, Michael Brammer, Philip M^c^Guire, Sagnik Bhattacharyya

## Abstract

High doses of delta-9-tetrahydrocannabinol (THC), the main psychoactive component of cannabis, have been shown to have anxiogenic effects. Also, THC effects have been shown to be modulated by genotype, including the single nucleotide polymorphism (SNP) rs1130233 at the protein kinase AKT1 gene, a key component of the dopamine signaling cascade. As such, it is likely that epigenetic methylation around this SNP may affect AKT gene expression, which may in turn impact on the acute effects of THC on brain function. We investigated the genetic (AKT1 rs1130233) and epigenetic modulation of brain function during fear processing in a 2-session, double-blind, cross-over, randomized placebo-controlled THC administration, in 36 healthy males. Fear processing was assessed using an emotion (fear processing) paradigm, under functional magnetic resonance imaging (fMRI). Complete genetic and fMRI data was available for 34 participants. THC caused an increase in anxiety and transient psychotomimetic symptoms and para-hippocampal gyrus/ amygdala activation. Number of A alleles at the AKT1 rs1130233 SNP, and percentage methylation at the CpG_11-12_ site, were independently associated with a greater effect of THC on activation in a network of brain regions including left and right parahippocampal gyri, respectively. AKT1 rs1130233 moderation of the THC effect on left parahippocampal activation persisted after covarying for methylation percentage, and was partially mediated in sections of the left parahippocampal gyrus/ hippocampus by methylation percentage. These results may offer an example of how genetic and epigenetic variations influence the psychotomimetic and neurofunctional effects of THC.

## 1. Introduction

Worldwide, cannabis is the most popular recreational drug (UNODC, 2019). Commonly smoked, but also ingested, there are over 140 different cannabinoids in cannabis (ElSohly, Radwan, Gul, Chandra, & Galal, 2017; Hanus, Meyer, Munoz, Taglialatela-Scafati, & Appendino, 2016; Pertwee, 2008b), with delta-9-tetrahydrocannabinol (THC) being the primary psychoactive cannabinoid (Pertwee, 2008b) responsible for its acute ‘high’ effects. Considerable variability in terms of sensitivity to the acute effects of cannabis has been found, with cannabis users reporting a wide range of subjective acute effects that are both positive, such as relaxation, happiness and laughter, as well as negative such as anxiety, panic attacks, and, less commonly, paranoia and psychosis-like symptoms (Ashton, 2001).

Consistent with preclinical evidence (Moreira & Wotjak, 2010), human experimental evidence suggests that such variability may be related to the specific dose of THC, with lower doses having anxiolytic (Phan et al., 2008) and higher doses having anxiogenic effects (Bhattacharyya, Atakan, et al., 2015; Bhattacharyya et al., 2017; Bhattacharyya, Fusar-Poli, et al., 2009; M. Colizzi et al., 2018). Further, the anxiogenic effects of THC seem to be attenuated when using cannabis strains containing also high-dose cannabidiol (CBD; (Boggs, Nguyen, Morgenson, Taffe, & Ranganathan, 2018)), the other main cannabinoid in cannabis. In line with this finding, under experimental conditions, CBD is known to have anxiolytic effects on its own (Bhattacharyya et al., 2010; Crippa et al., 2011), opposite neurophysiological and behavioral effects to those of THC when separately administered to the same individuals (Bhattacharyya, Crippa, et al., 2012b; Bhattacharyya, Falkenberg, et al., 2015; Bhattacharyya et al., 2010), and counteracting effects when co-administered along with THC (M. Colizzi & Bhattacharyya, 2017). Furthermore, we and others have provided evidence that sensitivity to the acute effects of THC on symptoms (Bhattacharyya, Crippa, et al., 2012a), cognition (Bhattacharyya et al., 2014) and their neurophysiological underpinnings (Bhattacharyya, Crippa, et al., 2012b) as well as to the short-term psychotomimetic effects of cannabis (van Winkel, Genetic, & Outcome of Psychosis, 2011), are moderated by a variation in the AKT1 gene (rs1130233). This gene codes for the protein kinase AKT, and its rs1130233 single nucleotide polymorphism (SNP) is a synonymous coding variation that has been linked to differential expression of the AKT protein, whereby the presence of an A allele is robustly associated with decreased expression of AKT (Blasi et al., 2011; Giovannetti et al., 2010; Harris et al., 2005; Tan et al., 2008).

The protein kinase AKT is a key component of the dopamine signaling cascade (Beaulieu, Gainetdinov, & Caron, 2007; Bozzi, Dunleavy, & Henshall, 2011) and altered AKT activity has been suggested to be relevant for the manifestation of psychiatric symptoms, including anxiety-like behaviors (Matsuda et al., 2019; Qiao et al., 2018). Regular cannabis use and acute THC administration have been shown to alter dopamine signaling in both human (Bossong et al., 2009; Sami, Rabiner, & Bhattacharyya, 2015) and animal (Bhattacharyya, Crippa, Martin-Santos, Winton-Brown, & Fusar-Poli, 2009) studies; and THC has been shown to modulate the phosphorylation of AKT1 (Ozaita, Puighermanal, & Maldonado, 2007; Shum et al., 2020). Altered expression of the AKT1 gene may therefore influence sensitivity to the effects of THC on brain functioning and related behavior, especially anxiety-related manifestations.

Methylation of DNA, i.e. addition of a methyl group to the cytosine pyrimidine ring of CpG dinucleotide, is one of several mechanisms of epigenetic control of gene expression, by reducing it (Suzuki & Bird, 2008). As such, methylation of the AKT1 gene is likely to reduce its gene expression and thereby indirectly reduce sensitivity to the acute effects of THC. Although we have previously demonstrated moderation of the acute effects of THC on functional brain activation by the AKT1 rs1130233 polymorphism (Bhattacharyya, Atakan, et al., 2012; Bhattacharyya et al., 2014), whether epigenetic interaction around this SNP also influences sensitivity to the acute effects of THC is yet to be investigated. Therefore, in the present study, we aimed to investigate the effect of an interaction between AKT1 rs1130233 polymorphism and methylation of CpG sites around this locus on the acute effect of THC administration on brain activation, during the processing of fear, as indexed using functional MRI. We focused on the effects of THC on fear processing-related brain activation in-light-of previous evidence that a single dose of THC modulates normal functioning of limbic regions involved in the processing of fear, in particular the amygdala, which correlated directly with the severity of anxiety induced by THC acutely (Bhattacharyya et al., 2017; Bhattacharyya et al., 2010). We hypothesized that the effect of a single dose of THC on fear-related activation of limbic regions would be modulated by both the AKT1 rs1130233 polymorphism and the methylation of CpG sites around this locus.

## 2. Methods

Ethical approval was granted for this study jointly buy the Institute of Psychiatry and South London and Maudsley NHS research committee and conducted in line with the declaration of Helsinki. Informed written consent was obtained from all participants before taking part in the study.

### Participants

Thirty-six right-handed healthy males, mean age 25.97± 5.8 years and IQ of 97.7 ± 6 took part in the study. None had a personal or family (first-degree relative) history of psychiatric illness. The Addiction Severity index was used to asses alcohol, cannabis and other illicit drug use (McLellan, Luborsky, Woody, & O’Brien, 1980)(Table 1). All had used cannabis at least once but no more than 25 times in their lifetime. None of them consumed illicit drugs regularly or over 21 units of alcohol per week. They were advised to not use illicit drugs, including cannabis for 30 days prior to the study or between the testing sessions. They were advised to have at least 8 hours sleep the night prior to each study day, and to abstain from consuming caffeine for 12 hours, tobacco for 4 hours, and alcohol for 24 hours prior to each study day. All subjects provided urine samples for drugs testing by immunometric assay kits. Of the subjects, 33 were white Europeans, two Sri-Lankan and one Chinese.

**Table 1:**
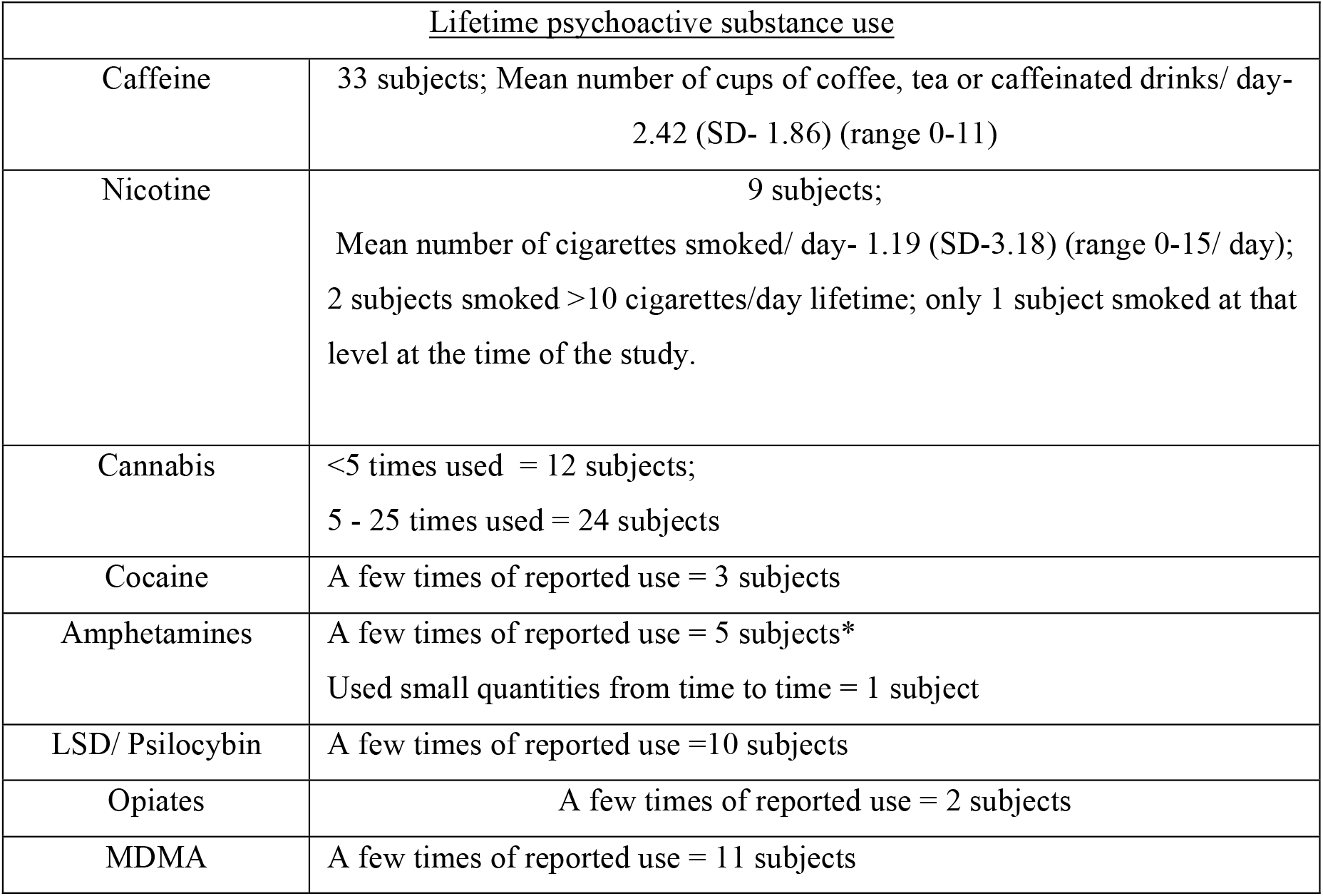
Previous use of psychoactive substances by particiapants (amended from our previous publication. (Bhattacharyya, Atakan, et al., 2015))

As genotyping at the AKT1 rs1130233 locus was unsuccessful in 1 participant, association between AKT1 rs1130233 SNP and methylation at CpG sites was investigated in 35 participants. Also, as another participant was unable to complete the fMRI scan under the THC condition, the effect of THC on activation compared to placebo and its correlation with percentage CpG methylation at site 11-12 were investigated in the 35 subjects. Thus, correlational analyses between genotype and THC effect on brain activation maps both with and without covarying for CpG methylation was performed in the 34 participants with completed neuroimaging and genotyping data.

### Experimental Design

As previously reported (Bhattacharyya, Atakan, et al., 2012; Bhattacharyya et al., 2014), each participant was tested on two separate occasions at least one month apart employing a double-blind, within-subject, crossover design. At each of the two sessions, they were given either 10mg of THC (approximately 99.6% pure, THC-pharm, Frankfurt, Germany) or a placebo (a matched gelatine capsule). To ensure that a roughly equal number of participants received THC or the placebo at each session, the order of drug administration was pseudo-randomised.

On the morning of each session, all participants passed a urine drug screening for opiates, cocaine, amphetamines, benzodiazepines and THC, using immunometric assay kits. Venous blood samples and psychopathological ratings were taken at baseline, and one, two and three hours post drug administration. A blood sample for genotyping and methylation analysis was obtained at baseline. Psychotomimetic effects were assessed by a clinician using the positive negative symptom scale (PANSS) (Kay, Fiszbein, & Opler, 1987). Spielberger state-trait anxiety inventory-state subscale (STAI-state) was used to measure self-rated anxiety (Spielberger, 1983). The analogue intoxication scale (AIS) was used to measure self-rated intoxication sedation (Mathew, Wilson, Humphreys, Lowe, & Wiethe, 1992), and the visual analogue mood scale was used to measure self-reported sedation (VAMS; the mental sedation subscale) (Norris, 1971). Subjects were scanned one hour after administration of THC or placebo. Pilot studies showed that the concentration of THC in blood samples plateaued and remained stable at approximately 1 to 2 hours after ingestion of the drug. We therefor peformed MRI scans from 1 hour after administration of the drug and the scans lasted for no more than 1 hour. During this scan participants completed the emotional processing task.

### Image acquisition

Images were acquired on a 1.5 Tesla Signa System (GE) at the Institute of Psychiatry, Psychology & Neuroscience, London. T2*-weighted images were taken using TR of 2000 msec; with 40msec echo time; a flip angle of 90° in 16 axial planes 7mm thick, parallel to anterior commissure posterior commissure line. To facilitate anatomical localisation of activation and high-resolution inversion recovery image dataset was acquired using a 3mm contiguous slices and an in-plane resolution of 3 mm (TR 16000 ms, Ti 180 ms, TE ms).

### fMRI task

Inside the scanner, participants performed the emotional (fear) processing task, which has been described in detail elsewhere (Bhattacharyya et al., 2017; Fusar-Poli et al., 2009). The blood oxygen level-dependant (BOLD) haemodynamic response was measured in subjects while they viewed faces that are neutral, mild or intensively fearful, or a crosshair. The subjects were asked to indicate the gender of the faces shown on the screen using a button-box. The faces were pseudo-randomised, each face was viewed for two seconds while the subjects were asked to press one of two buttons to determine the sex of the face. During the inter-stimulus interval, subjects were shown a fixation cross for 3-8 seconds according to a Poisson distribution. Performance data was collected in the form of reaction time and accuracy while indicating the gender of the facial stimuli.

### Genotyping and Methylation assay

DNA was extracted using standard methods by researchers at the Institute of Psychiatry (Freeman et al., 2003) and Genotyped for the AKT1 G>A rs1130233 SNP by KBioscience (Herts, UK; http://www.kbioscience.co.uk/) successfully for 35 out of the 36 subjects (corresponding to a call rate of 97%). Genotype frequencies and socio-demographic details of the 35 subjects are shown in Table 2. Genotype groups did not differ statistically significantly (p < .05) with regards to age, NART IQ or the number of years of education. Genotype frequencies for AKT1 at rs1130233 were in Hardy-Weinberg equilibrium (HWE; χ^2^=2.14, p>0.05) in the ethnically stratified sample.

**Table 2:** Participants by genotype, average age, NART IQ and education.

| Genotype | Age<br>(Mean±SD) | p | NART IQ<br>(Mean±SD) | p | Number of years of education<br>(Mean±SD) | p |
| --- | --- | --- | --- | --- | --- | --- |
| AKT1(G/G) (n=19) | 26.5±5.8 | NS | 98.8±6.9 | NS | 17.5±3.2 | NS |
| AKT1(G/A) (n=9) | 26.3±4.7 |  | 97.9±5.3 |  | 16.6±3.3 |  |
| AKT1(A/A) (n=7) | 23.8±6.6 |  | 95.5±8.0 |  | 17.0±7.4 |  |

The DNA methylation assay was carried out using previously described methods (Ouellet-Morin et al., 2013). Briefly, DNA samples were treated with sodium bisulfite using the EZ-96 DNA Methylation Kit (Zymo Research Corporation, Irvine, California) following the manufacturer’s protocol. Bisulfite–PCR amplification was conducted using Hot Star Taq DNA polymerase (Qiagen, UK). The primers used were GTTTTTGTTGAGTTAGGGTTTTTGA for the forward strand and TCCTTATCCAACATAAAATTCTCCA for the reverse strand, designed using the Sequenom EpiDesigner software (http://www.epidesigner.com). A total of 19 CpG sites were analysed in 14 CpG island blocks. This was such that CpG sites 6 and 7, 9 and 10, 11 and 12, 17 and 18, were analysed as a single site, due to their close proximity and high numbers of C and G nucleotides in between them, making them difficult to distinguish individually. Reactions were performed in duplicate and methylation analysis was carried out following established methods (Coolen, Statham, Gardiner-Garden, & Clark, 2007), using the Sequenom EpiTYPER platform (Sequenom Inc., USA), a reliable way of finding the density of methylated cytosines at specific genomic loci. Base-specific cleavage is followed by matrix-assisted laser desorption/ inonization-time of flight (MALDI-TOF) mass spectrometry. The size ratio of the cleaved products then provide methylation estimates for each cytosine-phosphate-guanine (CpG) unit, containing either one or an aggregate of neighbouring CpG sites. EpiTYPER software generated data that underwent stringent quality-control analysis where only CpG units with high calling rates (>80%) survived for statistical analyses. Genotyping and methylation assays were carried out blind to THC response results.

As a number of CpG sites are present around the rs1130233 locus, association between genotype and methylation status at each of these sites was examined first in order to identify the specific CpG sites to be further investigated in the present study. As there was a significant association between AKT1 rs1130233 genotype and methylation percentage at CpG sites 11-12 (CpG_11-12,_ Chr14: 104,773,527-104,773,522) around this locus (tested using a t-test), which survived correction for multiple testing (uncorrected *p*<0.001; corrected threshold *p*=0.0027), we focused on methylation at these sites for subsequent analyses.

### Image Analysis

The images were analyzed using a non-parametric software package, XBAM_v4.1, that was developed at the Institute of Psychiatry, Psychology & Neuroscience (King’s College London). It is important to use non-parametric measures when analyzing fMRI data as it may not follow a normal Gaussian distribution (Brammer et al., 1997; Thirion et al., 2007). Also, XBAM is less likely to misrepresent data distributions from outlier values as it uses medians rather than averages (Hayasaka & Nichols, 2003). The test statistic is calculated by standardizing for individual differences in residual noise before beging second-level, multi-subject testing, that uses a mixed effect-approach and robust permutation-based method.

Firstly, correction for head motion was completed by realigning the images, to a template created by computing a 3D volumes from the average intensity at each voxel throughout the whole period scanning (E. T. Bullmore, Brammer, et al., 1999). Realignment of the 3D image volume at each time-point to the template was computing using a combination of rotations (around the x, y and z axes) and translations (in x, y and z) that maximised the correlation between the template 3D volume and the image intensities of the volume in question. In order to smooth the data a 7.2mm full-width-at-half-maximum Gaussian filter was applied to average the relative intensities of neighbouring voxels. Indervidual activation maps were created by modelling the BOLD signal using 2 gamma-variate functions for each experimental condition, peaking at 4 and 8 seconds to allow for variability in haemodynamic delay. A best fit between the weighted sum of these convolutions and the change over time was computed at each voxel using the constrained BOLD effects model (Friman, Borga, Lundberg, & Knutsson, 2003). This method increases the robustness of the model fitting procedure to not give mathematically plausible, but physiologically implausible results. The sum of squares (SSQ) ratio (calculated as the ratio of the SSQ of deviations from the mean image intensity due to the model component over the whole time series to the SSQ of deviations due to the residuals) was estimated for each voxel, followed by permutation testing, determining which voxels were significantly activated(E. Bullmore et al., 2001). This avoids F statistic associated problems, where in fMRI time series the use of the residual degrees of freedom are often unknown due to coloured noise in the signal. Data was then permuted by the wavelet-based method; both described and characterized previously (E. Bullmore et al., 2001), this permits a data driven calculation of the null distribution of SSQ, vy assuming no experimentally-determined response. This distribution was then used to threshold the activation maps at the desired Type 1 error rate of less than one false positive. Indervidual SSQ ratio maps were transformed into standard stereotactic space (Talairach & Tournoux, 1988) using a two-stage warping procedure (Brammer et al., 1997) allowing for localization of activations. Initially an individual average image intensity map over the course of the experiment was computed, followed by computation of the affine transformations required to map this image to first their structural scan and then to the Talairach template, by maximizing between image correlations at each stage. The BOLD effect size and SSQ ratio maps were transformed into Talairach space using these transformation methods.

Group activation maps were created for each task condition (‘intensely fearful faces’, ‘mildly fearful faces’, ‘neutral faces’), and for each drug condition (THC, placebo). Median SSQ ratios at each voxel across all individuals were determined in the observed and permutated data maps, using medians to minimize outlier effects. A null distribution of SSQ ratios was driven from the distribution of median SSQ ratios over all intracerebral voxels from the permuted data, giving group activation maps for each condition. We could then directly compare group activation at each condition using non-parametric repeated-measure analysis of variance (ANOVA) (Brammer et al., 1997).

Activated voxels were grouped into clusters of activation using a method previously described (E. T. Bullmore, Suckling, et al., 1999), and shown to give excellent cluster-wise Type 1 error control. A voxel-wise statistical threshold of p=0.05 was used and the cluster-wise thresholds were choosen such that the number of false positive clusters would be <1 per brain (therefor we have only reported regions that survive both the critical statistical threshold and the corresponding p values from cluster-level analysis). The data from more than one voxel is integrated into the test statistic giving greater sensitivity and allows for a reduction in the search volume or overall number of required tests for whole brain analysis, thereby helping to mitigate the problem multiple comparisons. For each drug condition, we therefore had a separate standard-space map for each of the experimental conditions (‘intensely fearful faces’, ‘mildly fearful faces’ or ‘neutral faces’) for each participant. We then employed a non-parametric repeated measures analysis of variance (ANOVA) whole-brain analysis approach to identify brain regions that were activated by THC relative to the placebo condition. We contrasting the individual brain activation maps for intense and mildly fearful faces combined (as the fear condition) with those of the neutral faces.

As we were interested in investigating the relationship of AKT1 rs130233 genotype and methylation (at CpG site 11-12 around that locus) with the cerebral activation response to THC during fear processing, we followed the Baron and Kenney (Baron & Kenny, 1986)mediation analysis approach to examine whether methylation at CpG site 11-12 (moderator variable) mediated the relationship between brain activation during fear processing (dependent variable) and AKT1 SNP rs1130233 (independent variable). Therefore we investigated the association between independent variable (genotype) and dependent variable (brain activation), then between moderator variable (methylation) and dependent variable and between moderator variable and independent variable by carrying out separate correlational analyses. Finally, to investigate the extent to which the association between genotype and THC effect on fear-related brain activation was mediated by methylation at the CpG 11-12 site, we carried out a final correlational analysis between genotype (AKT rs130233) and the effect of THC on brain activation during fear processing, while covarying for methylation percentage at the CpG site 11-12.

These analysis examined the association between the median SSQ ratio under each drug condition (THC and placebo) while processing fear with : 1) genotype (“number of A alleles +1”, at the AKT1 rs1130233 locus, coded as G/G =1 G/A=2 and A/A= 3); and 2) DNA methylation (methylation percentage at CpG site 11-12). We contrasted each of the active (‘intensely fearful faces’ and ‘mildly fearful faces’) conditions of the fear processing task with the baseline condition (‘neutral faces’) to generate contrast of interest maps (‘intense fear’ map: ‘intensely fearful minus neutral face’; ‘mild fear’ map: ‘mildly fearful minus neutral face’) for each individual participant under each drug condition. We estimated the Pearson’s product moment correlation coefficient at each voxel, in each subject’s standard space, yielding one correlation coefficient (r) per intracerebral voxel. Correlation differences between groups were estimated at each voxel by computing for each group, the **r** value for each subject followed by subtracting the resulting two values. A null distribution was then appropriately generated by randomly permuting subjects and their associated genotype or methylation levels between the groups (without replacement), therefore scrambling any group differences. For each permutation, the difference in correlation between the scrambled groups was calculated and then the resulting values combined for all voxels to produce a whole-brain null distribution of differences in correlation. The critical value for significance at any desired Type 1 error level in the original (non-permuted) data was then obtained from this distribution after sorting it and selecting the appropriate point from the sorted distribution. This means that the critical value for a one-tailed test thresholded at p=0.05 would be the value of the difference in correlation in the null distribution where 95% of all the null values lay below that point. The statistical analysis was then extended to the 3D cluster level, previously described (E. T. Bullmore, Suckling, et al., 1999). The probability for each cluster was chosen to set the level of expected Type I error clusters to less than 1 error cluster per whole brain under the null hypothesis.

## 3. Results

### Participants

All subjects had used cannabis at least once, but no more than 25 times in there lifetime. No participant consumed more than 21 units of alcohol per week, and all had low levels of illicit drug use. None of the participants had used cannabis or other illicit drugs for at least one month prior to their first visit, and they were asked to abstain from use between visits. Substances use information in this cohort of participants has been reported previously (Bhattacharyya, Atakan, et al., 2015). It has been presented again in this manuscript in the form of an amended table (Table 1) for the sake of completeness.

Genotype frequencies of participants showed no difference between groups with regard to age, NART IQ and number of years of education, shown in Table 2 for all 35 participants successfully genotyped. Genotype data for both genetic variants were tested for deviation from Hardy-Weinberg equilibrium (HWE). Frequencies for AKT rs1130233 showed a deviation from the HWE (χ^2^=6.0, p<0.05). However, the AKT1 rs1130233 polymorphism was retained consistent with current practice, as there was no evidence of a quality control issue and also as recent evidence suggests that while polymorphisms that are not in HWE may be less powerful, they do not tend to increase false positive results (Fardo, Becker, Bertram, Tanzi, & Lange, 2009).

### Symptomatic and behavioral response to THC

These results have been previously reported in detail by Bhattacharyya et al. (2015), however, we are summarising them here again for completeness. As we have previously reported (Bhattacharyya, Atakan, et al., 2015), administration of THC was associated with change in psychopathological ratings over time (estimated as the area under the curve; AUC). In particular, THC administration was associated with the induction of transient positive psychotic symptoms (as indexed using the PANSS positive symptoms subscale; THC (AUC) = 25.83 ± 6.0, Placebo (AUC) = 21.96 ±1.8, p<0.001)) and anxiety symptoms (as indexed using the STAI-state, THC (AUC)= 49.36 ± 27.6, PLB (AUC)= 37.12 ±22.3, p<0.001)). Also, a THC-induced change in total PANSS scores (THC (AUC)= 108.61 ± 16.6, PLB (AUC)= 94.49 ± 6.4, p<0.001) and levels of sedation (as indexed using the VAMS, THC (AUC) = 54.98 ± 14.8, PLB (AUC)= 48.65 ± 16.2, p=0.02; shown in our previous publication (Bhattacharyya, Atakan, et al., 2015)) and intoxication (as indexed using the AIS, THC (AUC)= 9.99 ± 5.5, PLB (AUC)= 3.64 ± 4.4, p<0.001) were observed.

There was no significant effect of drug condition on reaction time during the emotional (fear) processing task (p=0.597) or performance accuracy across all emotional conditions (p=0.933) as well as for fear (p=0.976). There was no statistically significant effect of drug condition on reaction time during the emotional (fear) processing task (p=0.597) or performance accuracy across all emotion conditions (p=0.933) as well as for fear (p=0.976).

### Regional brain response to THC during fear processing

There was a statistically significant effect of THC administration on the normal pattern of regional brain activation associated with fear in a distributed network of regions including the left parahippocampal gyrus and amydgala. THC enhanced left parahippocampal gyrus/amydgala engagement while viewing fearful faces compared to neutral faces, while there was an attenuation of engagement in these regions under placebo while viewing fearful faces relative to neutral faces (Figure 1).

**Figure 1.**
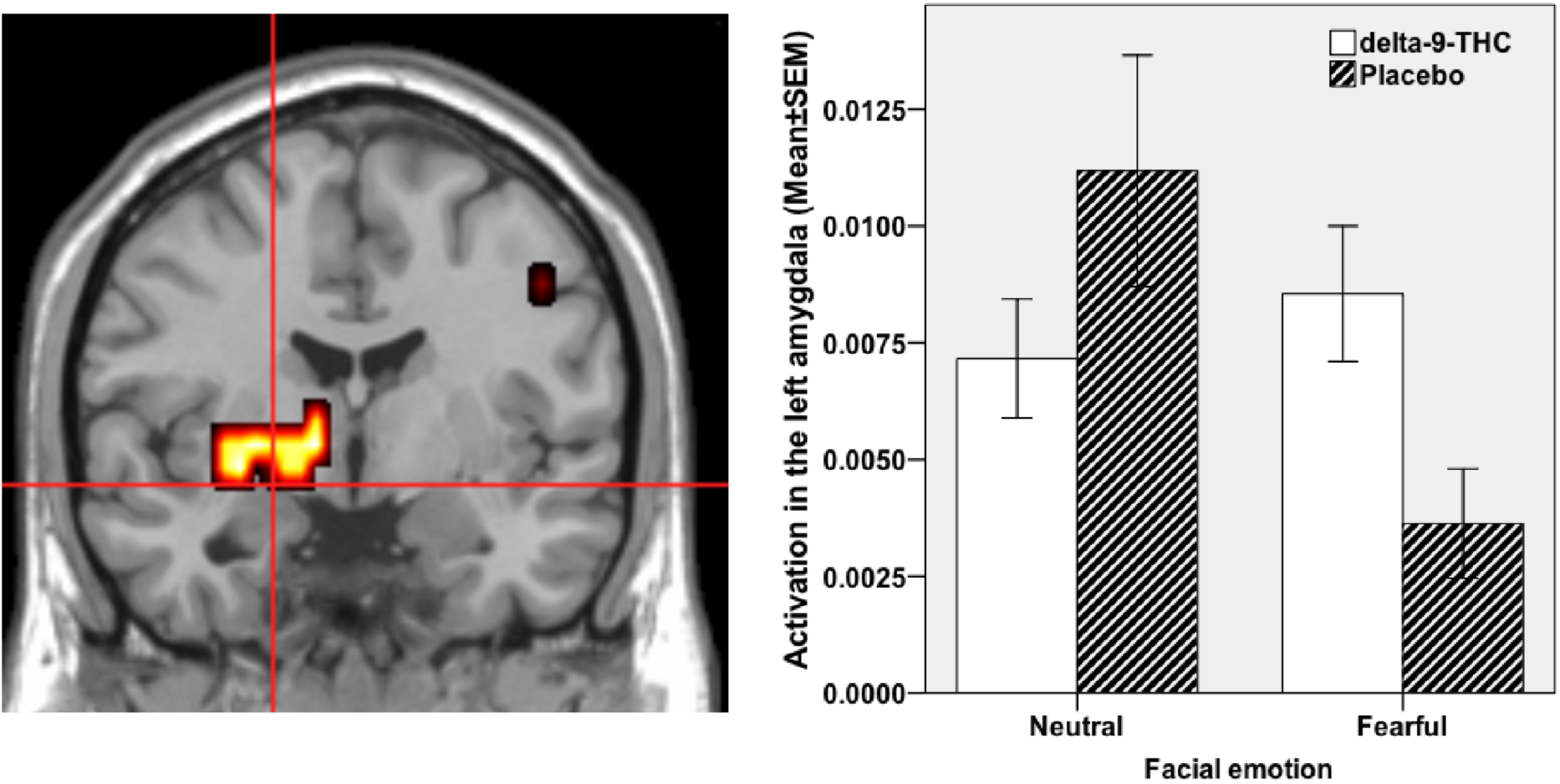
Brain activation whilst viewing fearful faces in comparison to neutral faces under THC condition compared to Placebo. Activation shown in the left parahippocampal gyrus and amygdala (cross-hairs in the coronal view; cluster size= 18 voxels; Talairach coordinates: *x*= -29, *y*= -4, *z*= -7; *x*=-29, *y*=7, *z*=-13; *p*<0.003 corrected for <1 false positive cluster). The left side of the brain is shown on the left side of the image. Accompanying plot on the right shows activation in the amygdala under the different fear (fearful vs neutral faces) and drug (THC vs PLB) conditions.

### Relationship between AKT1 genotype (rs1130233) and the effect of THC on regional brain activation during fear processing

There was a statistically significant association between the effect of THC on regional brain activation while processing fear and AKT1 genotype (rs1130233), such that the higher the number of risk alleles (A) the greater was the effect of THC on fear-related brain activation across a network of brain regions (Figure 2 A and Table 3) that included parahippocampal, fusiform and cingulate gyri. Additionally, a cluster in the left anterior cingulate gyrus/ medial prefrontal cortex (x= -11, y= 26, z= 26; rho=0.395, p=0.021) correlated positively with change in STAI scores.

**Figure 2.**
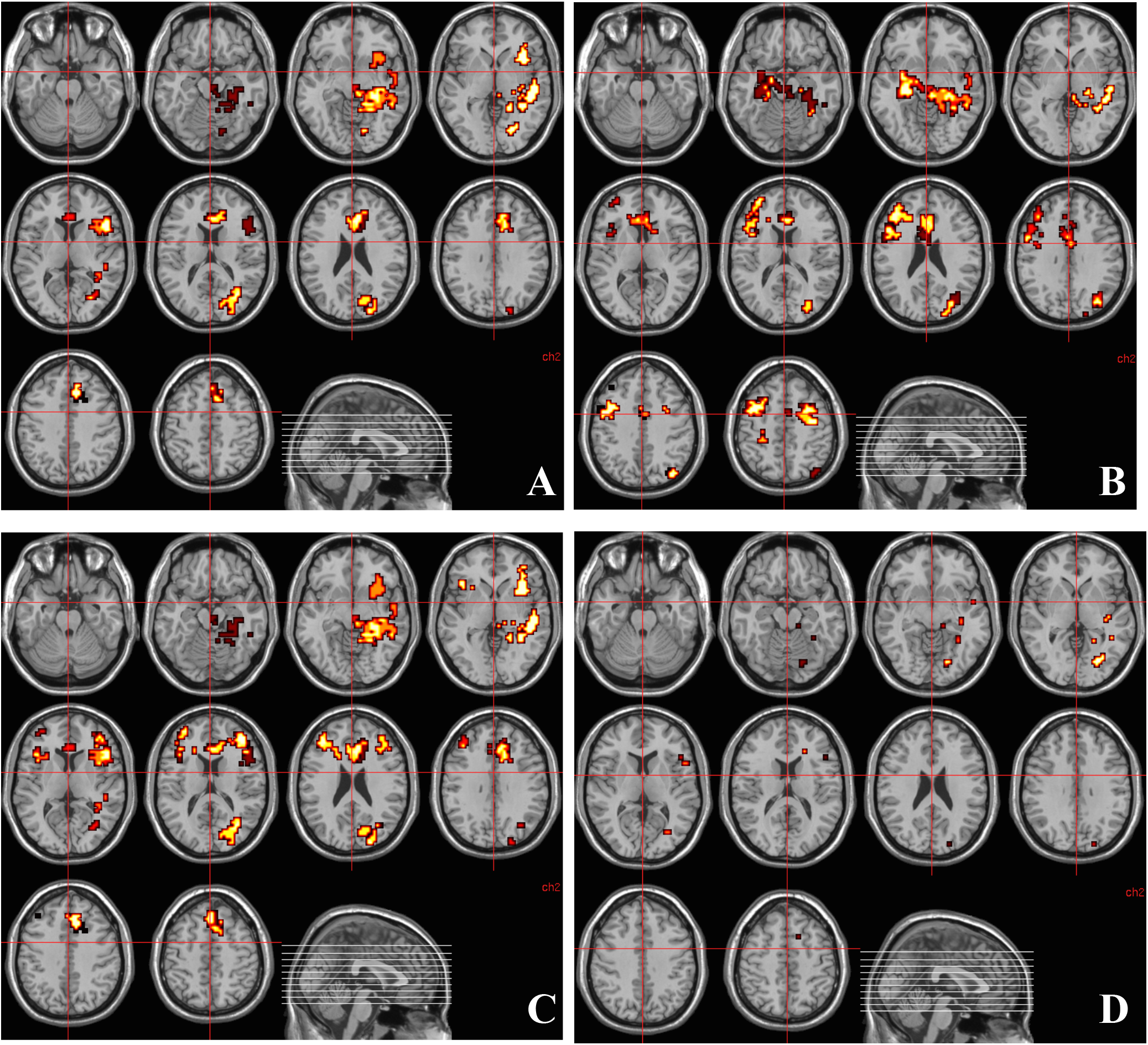
A - Regions of brain activation showing significantly greater correlation with the number of A alleles of the AKT1 rs1130233 polymorphism under THC condition than under placebo condition while viewing fearful faces. The left side of the brain is shown on the left side of the image. B - Regions of brain activation showing significantly greater correlation with methylation percentage at CpG island 11-12 around rs1130233 polymorphism in the AKT1 gene under THC condition than under placebo condition while viewing fearful faces. The left side of the brain is shown on the left side of the image. C-Regions of brain activation showing significantly greater correlation with the number of A alleles of the AKT1 rs1130233 polymorphism under THC condition than under placebo condition while viewing fearful faces after controlling for percentage methylation at CpG island 11-12 around rs1130233 polymorphism in the AKT1 gene. The left side of the brain is shown on the left side of the image. D - Clusters of brain activation where the significantly greater correlation between the number of A alleles of the AKT1 rs1130233 polymorphism under THC condition than under placebo condition while viewing fearful faces may be mediated by correlation with methylation percentage at CpG island 11-12 around rs1130233 polymorphism in the AKT1 gene. The left side of the brain is shown on the left side of the image.

**Table 3.** Regions of brain activation (Talairach coordinates) showing significantly greater correlation with the number of A alleles of the AKT1 rs1130233 polymorphism under THC condition than under placebo condition while viewing fearful faces. L and R indicate the left and right hemisphere.

| Cerebral Region | x | y | z | Size (no. of voxels) | Side | p-Value |
| --- | --- | --- | --- | --- | --- | --- |
| Inferior Frontal Gyrus | -43 | 26 | 4 | 14 | L | 0.006583 |
|  | -40 | 26 | -2 | 22 | L |  |
|  | -32 | 22 | -13 | 15 | L |  |
|  | -36 | 19 | -7 | 17 | L |  |
| Anterior Cingulate Gyrus | -4 | 22 | 37 | 10 | L | 0.002992 |
|  | -11 | 30 | 20 | 15 | L |  |
| Fusiform Gyrus. | -22 | -67 | -7 | 8 | L | 0.002992 |
| Cuneus. | -11 | -74 | 15 | 10 | L | 0.002992 |
| Middle Occipital Gyrus. | -29 | -63 | 4 | 5 | L | 0.002992 |
|  | -25 | -81 | 9 | 25 | L |  |
|  | -18 | -85 | 15 | 8 | L |  |
| Culmen. | -18 | -44 | -7 | 5 | L | 0.001496 |
|  | -22 | -44 | -18 | 7 | L | 0.008378 |
| Middle Temporal Gyrus | -51 | -33 | -13 | 63 | L | 0.001496 |
|  | -51 | -11 | -13 | 7 | L |  |
| Brainstem, Midbrain, | 0 | -15 | -18 | 5 | R | 0.001496 |
| Parahippocampal gyrus | -40 | -30 | -7 | 30 | L | 0.001496 |
|  | -25 | -26 | -18 | 12 | L |  |
| Cingulate Gyrus. | -7 | 19 | 42 | 5 | L | 0.002992 |
|  | -11 | 26 | 26 | 14 | L |  |
|  | -11 | 26 | 31 | 10 | L |  |
| Anterior Cingulate. | 0 | 30 | 15 | 20 | R | 0.002992 |
|  | -7 | 33 | 9 | 13 | L |  |
|  | 0 | 37 | 4 | 7 | R |  |

### Relationship between methylation at CpG_11-12_ site around AKT1 SNP (rs1130233) and the effect of THC on regional brain activation during fear processing

There was a significant association between the effect of THC on regional brain activation while processing fear, and methylation at CpG_11-12_ site, such that the higher the percentage of methylation at this site, the greater was the effect of THC on fear-related brain activation across a network of brain regions (Figure 2 B and Table 4) that included parahippocampal and cingulate gyri. Anxiety induced under the influence of THC (indexed using STAI) correlated positively with its effect on parahippocampal activation (x=29, y=-22, z=-13; rho= 0.401, p=0.017).

**Table 4.** Regions of brain activation (Talairach coordinates) showing significantly greater correlation with methylation percentage at CpG island 11-12 around rs1130233 polymorphism in the AKT1 gene under THC condition than under placebo condition while viewing fearful faces. L and R indicate the left and right hemisphere.

| Cerebral Region | x | y | z | Size (no. of voxels) | Side | p-Value |
| --- | --- | --- | --- | --- | --- | --- |
| Superior Frontal Gyrus | 36 | 52 | 15 | 27 | R | 0.003249 |
| Medial Frontal Gyrus | 40 | 48 | 9 | 11 | R | 0.003249 |
|  | 43 | 37 | 20 | 9 | R |  |
|  | 43 | 7 | 42 | 20 | R | 0.002067 |
|  | 40 | 11 | 37 | 26 | R |  |
|  | 40 | 7 | 48 | 15 | R |  |
|  | 0 | -4 | 53 | 9 | R | 0.005907 |
| Inferior Frontal Gyrus | 51 | 26 | 20 | 12 | R | 0.003249 |
|  | 43 | 4 | 31 | 17 | R | 0.002067 |
| Sub-Gyral | 40 | 15 | 15 | 12 | R | 0.003249 |
|  | 11 | 30 | -2 | 8 | R | 0.002658 |
|  | -43 | -33 | -7 | 18 | L |  |
|  | -47 | -15 | -13 | 5 | L |  |
|  | -25 | 7 | 42 | 26 | L | 0.005907 |
|  | -29 | 7 | 37 | 10 | L |  |
|  | -32 | -4 | 37 | 6 | L |  |
|  | 25 | -4 | 53 | 23 | R | 0.007383 |
|  | -29 | -70 | 26 | 5 | L | 0.009155 |
| Middle Occipital Gyrus | -25 | -78 | 15 | 9 | L | 0.009155 |
|  | -25 | -74 | 9 | 8 | L |  |
| Middle Temporal Gyrus. | -29 | -70 | 20 | 11 | L | 0.009155 |
| Precentral Gyrus. | 29 | -15 | 48 | 21 | R | 0.007383 |
| Precuneus. | -36 | -70 | 37 | 5 | L | 0.009155 |
| Angular Gyrus. | -36 | -74 | 31 | 6 | L | 0.009155 |
| Fusiform Gyrus. | -32 | -37 | -18 | 18 | L | 0.002658 |
| Midbrain | -4 | -26 | -13 | 47 | L | 0.002658 |
| Left Brainstem, Midbrain | -4 | -22 | -18 | 6 | L | 0.002658 |
| Insula | 40 | 15 | 9 | 13 | R | 0.003249 |
| Extra-Nuclear Corpus<br>Callosum. | 4 | 15 | 20 | 12 | R | 0.002658 |
|  | -4 | 26 | -2 | 8 | L |  |
| Amygdala/ Parahippocampal<br>gyrus | 22 | -7 | -18 | 36 | R | 0.003249 |
| Parahippocampal Gyrus. | 29 | -22 | -13 | 23 | R | 0.003249 |
| Anterior Cingulate. | 0 | 37 | 4 | 9 | R | 0.002658 |
|  | 4 | 33 | 9 | 6 | R |  |
|  | 0 | 30 | 15 | 19 | R |  |

### Relationship between methylation at CpG_11-12_ site around AKT1 SNP (rs1130233) and the number of A alleles at AKT1 SNP (rs1130233)

As expected, there was a linear relationship (r=0.869, p=<0.001) between the number of A alleles at AKT1 rs1130233 and methylation percentage at CpG sites 11-12, (therefore the higher the number of A alleles the higher the methylation percentage).

### Relationship between AKT1 genotype (rs1130233) and the effect of THC on regional brain activation during fear processing after covarying for methylation at CpG_11-12_ site

On investigating whether the association between AKT1 rs1130233 genotype and THC-induced change in brain activation while processing fear was mediated by degree of methylation at the CpG_11-12_ site, we found that some of the association between AKT1 rs1130233 polymorphism and effect of THC on fear-related brain activation (as depicted in Figure 3.A and table 1) was no longer present (as depicted in Figure 2 C and Table 5) after covarying for percentage methylation at the CpG_11-12_ site. This suggested that methylation percentage at the CpG_11-12_ site partly mediated some of the association between AKT1 rs1130233 polymorphism and the effect of THC on fear-related brain activation. Brain regions where the association between AKT1 rs1130233 genotype and THC’s effect on fear-related activation were mediated by percentage methylation at the CpG_11-12_ site around AKT1 rs1130233 localize to the left parahippocampal gyrus extending toward the lingual gyrus, left fusiform gyrus, midbrain extending to the left parahippocampal gyrus, left hippocampus extending towards the insula and the left superior temporal gyrus (Figure 2 D and Table 6).

**Table 5.** Regions of brain activation (Talairach coordinates) showing significantly greater correlation with the number of A alleles of the AKT1 rs1130233 polymorphism under THC condition than under placebo condition while viewing fearful faces after controlling for percentage methylation at CpG island 11-12 around rs1130233 polymorphism in the AKT1 gene. L and R indicate the left and right hemisphere.

| Cerebral Region | x | y | z | Size (no. of voxels) | Side | p-Value |
| --- | --- | --- | --- | --- | --- | --- |
| Middle Frontal Gyrus. | 40 | 52 | 4 | 3 | R | 0.005075 |
|  | 36 | 48 | 9 | 7 | R |  |
|  | 36 | 48 | 15 | 21 | R |  |
|  | -36 | 44 | 9 | 17 | L |  |
|  | -40 | 41 | 15 | 13 | L |  |
|  | 40 | 37 | 20 | 8 | R |  |
| Medial Frontal Gyrus. | 0 | 33 | 42 | 9 | R | 0.005075 |
|  | 0 | 30 | 37 | 15 | R |  |
| Inferior Frontal Gyrus. | 40 | 26 | 4 | 7 | R | 0.005075 |
|  | 40 | 26 | -7 | 5 | R |  |
|  | -47 | 19 | 4 | 13 | L |  |
| Middle Temporal Gyrus. | -51 | -11 | -13 | 7 | L | 0.007761 |
|  | -51 | -33 | -13 | 63 | L |  |
| Sub-Gyral. | -32 | 44 | 4 | 12 | L | 0.005075 |
|  | -40 | -30 | -7 | 28 | L | 0.007761 |
|  | -36 | -37 | -2 | 4 | L |  |
| Middle Occipital Gyrus. | -22 | -85 | 15 | 9 | L | 0.007761 |
|  | -29 | -63 | 4 | 5 | L |  |
| Culmen. | -18 | -44 | -7 | 4 | L | 0.007761 |
| Cuneus. | -11 | -74 | 15 | 10 | L | 0.007761 |
|  | -18 | -85 | 20 | 3 | L |  |
| Extra-Nuclear. | -32 | 22 | -2 | 25 | L |  |
|  | 32 | 22 | -2 | 7 | R | 0.005075 |
|  | -32 | 15 | -7 | 25 | L |  |
| Left Brainstem, Midbrain. | 0 | -15 | -18 | 4 | R | 0.007761 |
| Parahippocampal Gyrus. | -25 | -26 | -18 | 18 | L | 0.007761 |
| Posterior Cingulate. | -22 | -67 | 9 | 27 | L | 0.007761 |
| Cingulate Gyrus. | -7 | 19 | 42 | 7 | L | 0.005075 |
|  | -11 | 26 | 26 | 15 | L |  |
|  | -11 | 26 | 31 | 14 | L |  |
| Anterior Cingulate. | -11 | 26 | 20 | 17 | L | 0.005075 |
|  | 0 | 30 | 15 | 22 | R |  |
|  | -4 | 33 | 9 | 12 | L |  |
|  | 0 | 37 | 4 | 7 | R |  |

**Table 6.**
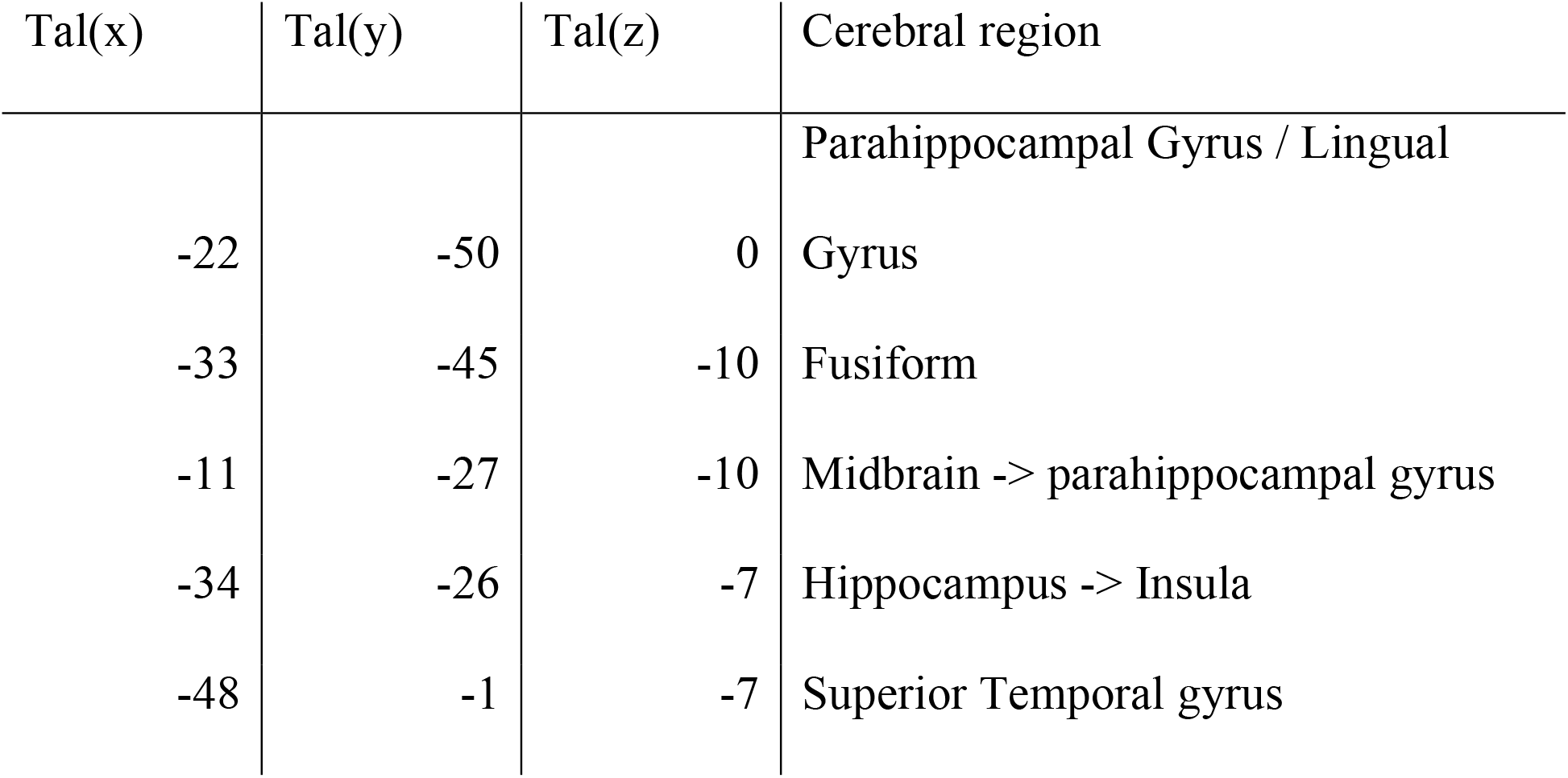
Clusters of brain activation where the significantly greater correlation between the number of A alleles of the AKT1 rs1130233 polymorphism under THC condition than under placebo condition while viewing fearful faces may be mediated by correlation with methylation percentage at CpG island 11-12 around rs1130233 polymorphism in the AKT1 gene. L and R indicate the left and right hemisphere.

## 4. Discussion

Here we investigate the independent effects of AKT1 SNP rs1130233 genotype and methylation levels at its surrounding CpG sites as well as their relationship with fear-related brain activity and associated anxiety-like behavior in healthy individuals while they were under the acute influence of experimentally administered THC. We found an association between genotype alone and THC effect on brain activation, such that, following acute THC exposure allele A (previously shown to moderate sensitivity to the acute psychoactive effects of THC (Bhattacharyya, Atakan, et al., 2012; Bhattacharyya et al., 2014)) was associated with greater activation in the parahipocampal and cingulate gyri during fear processing. Similarly, we found that, following acute THC exposure, increased methylation percentage around the SNP locus was associated with greater activation in the same brain regions whilst fear processing. Also, the greater the activation of such brain regions, the more severe were the anxiety symptoms following acute THC administration. Finally, the most novel finding from this study is that, following acute THC exposure, methylation around the rs1130233 SNP partially mediated the effect of AKT1 genotype on fear processing in the parahippocampal gyrus and hippocampus, extending to the STG and insula.

Convergent evidence from both animal (Lai et al., 2006) and human (Tan et al., 2008) studies implicates AKT1 in modulating prefrontal-striatal structure and function and suggests that its deficiency creates a context permissive for gene-gene and gene-environment interactions that contribute to altered dopaminergic transmission, increasing the risk of dopamine-associated disorders and behaviors. Consistent with this, several previous studies have indicated that genetic variations in the AKT1 gene may moderate the acute psychotogenic effects of cannabis (Bhattacharyya, Atakan, et al., 2012; Bhattacharyya et al., 2014; Lai et al., 2006; Morgan, Freeman, Powell, & Curran, 2016; Tan et al., 2008), although this has not been confirmed by one recent study (Hindocha et al., 2020), possibly due to differences in study design. The present study extends such findings indicating that the rs1130233 polymorphism of the AKT1 gene also moderates the acute effects of THC on the neurophysiological underpinnings of fear processing and associated anxiety-like behavior.

Previous work has explored how continued exposure to cannabis and its main psychoactive compound THC may result in aberrant epigenetic modifications, including altered methylation (Szutorisz & Hurd, 2016). However, limited investigation has been carried out so far on how epigenetic mechanisms may modulate behavioral responses to cannabis, including psychotomimetic symptoms. While increased methylation from another dopamine-related gene, catechol-O-Methyltransferase (COMT), has been associated with lower cannabis use frequency during adolescence (van der Knaap et al., 2014), the present study is the first to investigate the effects of methylation in the AKT1 gene on brain functioning and related behavior following acute THC exposure. Specifically, under the effects of THC, we found greater functional activation in fear processing-related brain regions and more severe associated anxiety, as a function of increasing AKT1 methylation. Methylation levels also partially mediated the modulatory effect of genetic variation in the AKT1 gene on acute THC-induced activation of brain regions involved in fear processing.

While viewing neutral faces, acute THC exposure decreased brain activation consistent with the known effects of cannabis (Pertwee, 2008a) on brain activation (Batalla et al., 2014; Bossong, Jager, Bhattacharyya, & Allen, 2014). Instead, acute THC administration increased activation when viewing fearful faces in limbic regions such as the parahippocampal gyrus and amygdala. As such increase in fear processing-related brain activation was associated with both the load of AKT1 rs1130233 allele A and related methylation (and thus lower gene expression), then altered AKT1 activity may therefore represent a marker for differing responses to fear processing under the acute effects of cannabis. More specifically, the THC-induced increase in brain activation in several brain regions involved in emotional processing as a function of both these genetic factors may possibly represent a genetically mediated lack of efficiency in processing emotion, similar to that found by previous studies (Lai et al., 2006; Tan et al., 2008). Such explanation would be corroborated by the evidence that two clusters in the cingulate gyrus and inferior frontal gyrus also positively correlated with the STAI score, an indication that increased activation in these brain regions may reflect higher levels of anxiety.

Presence of the A allele at this SNP has been associated with lower AKT1 expression (Blasi et al., 2011; Giovannetti et al., 2010; Harris et al., 2005; Tan et al., 2008) and increased methylation levels usually do also often have the effect of decreasing gene expression (Suzuki & Bird, 2008). As both the rs1130233 A allele and increased methylation around this locus were associated with THC-induced increased activation in a number of brain regions, such neurophysiological effects of the drug could be due to the effect of both genotype and methylation in reducing AKT1 gene expression (Emamian, Hall, Birnbaum, Karayiorgou, & Gogos, 2004). Acute THC exposure has been shown to result in an increase in dopamine release (Bloomfield, Ashok, Volkow, & Howes, 2016) via a CB1 receptor-dependent modulation of glutamate (M. Colizzi & Bhattacharyya, 2018; M. Colizzi, McGuire, Pertwee, & Bhattacharyya, 2016; M. Colizzi et al., 2019) and GABA signaling (Laruelle & Abi-Dargham, 1999), leading to the manifestation of psychotomimetic symptoms (M. Colizzi et al., 2019). Dopamine binding at its receptors has an inhibitory effect on AKT1 action (Beaulieu, Tirotta, et al., 2007) and THC has also been reported to induce the phosphorylation of AKT1 (Sánchez, Ruiz-Llorente, Sánchez, & Díaz-Laviada, 2003), reducing AKT action even beyond acute intoxication. THC-induced dopamine release may therefore be increased in the context of genetically-determined lower AKT1 gene expression, because of reduced feedback regulation in the cascade where the AKT protein is present, as well as because of a direct action of THC in reducing the expression of the AKT protein (Ozaita et al., 2007).

Differing neurophysiological and behavioral effects of acute THC exposure, dependent on AKT1 genetic variation, with mediation via methylation levels, may begin to offer examples for specific genetic markers that are associated with an increased risk for neuropsychiatric disorders following cannabis use. AKT1 signaling cascade affects dopamine 2 (D2) receptors, encoded by the DRD2 gene, known to play a key role in psychosis (Wong et al., 1986), and decreased AKT1 expression could result in an increased level of synaptic activation through the inhibitory effects from dopamine binding on the enzyme (Arguello & Gogos, 2008). Consistent with this, another SNP in the AKT1 gene, rs2494732, which is in high linkage disequilibrium with rs1130233 (*r*^2^=0.45, *D*′=0.94), entails a variant – allele A – which has been shown to increase the risk of developing psychosis among cannabis users (Di Forti et al., 2012), and to interact with genetic variation in the DRD2 gene in further increasing the risk of psychosis possibly because of genetically-determined higher striatal dopamine levels (M. Colizzi et al., 2015).

By covarying for methylation levels around the genotype locus, the present findings can only be used to make suggestive comments about the effect of methylation. Therefore, replication of our results is required to gain stronger evidence into the independent effect of methylation and to further investigate the specific brain regions potentially involved. Also, the results from this study should only be considered as preliminary due to the small number of participants. Furthermore, the relationship between peripheral epigenetic markers and brain levels is still uncertain. Future studies should try to examine methylation as a possible attenuation mechanism for gene expression, regardless of genotype and alongside genotype, and how it mediates the effects of acute THC exposure at the molecular level. Other SNPs in the AKT1 gene also warrant further research to identify their role in gene expression and how they modulate THC response. This study is further limited using a 1.5 Tesla scanner; as more powerful scanners has now become more routinely available.

In conclusion, this work further adds to existing evidence for a role of the AKT1 SNP rs1130233 in modulating the acute effects of THC during fear processing, with these being associated with the A allele presence. Also, we have provided the first suggestive evidence that methylation around the SNP, may also independently modulate such acute response to THC. Finally, methylation levels mediated the effect of the AKT1 SNP rs1130233 in increasing brain activation following acute THC exposure whilst processing fear, suggesting a mechanism of further divergence in individuals’ anxiogenic responses to cannabis.

## Acknowledgements, conflicts of interests and ethical standards

This work was supported by a Joint Medical Research Council/Priory Clinical research training fellowship (G0501775) from the Medical Research Council, United Kingdom, to SB. SB has also been supported during this work by the National Institute for Health Research (NIHR), UK through a Clinician Scientist award (NIHR-CS-11-001), a grant from the NIHR Efficacy and Mechanism Evaluation scheme and by the NIHR Biomedical Research Centre for Mental Health at the South London and Maudsley NHS Foundation Trust and Institute of Psychiatry, Psychology and Neuroscience, King’s College London jointly funded by the Guy’s and St Thomas’ Trustees and the South London and Maudsley Trustees. DP was supported, during this work, by the European Commission Seventh Framework Programme Marie Curie Career Integration Grant FP7-PEOPLE-2013-CIG-631952, the 2016 Bial Foundation Psychophysiology Grant grant - Ref. 292/16, and the Fundação para a Ciência e Tecnologia (FCT) IF/00787/2014, LISBOA-01-0145-FEDER-030907 and DSAIPA/DS/0065/2018 grants, and the iMM Lisboa Director’s Fund Breakthrough Idea Grant 2016. There are no known conflicts of interest associated with this publication and no significant financial support for this work that could have influenced any of the authors other than DP who is a co-founder and shareholder of the neuroimaging research services company NeuroPsyAI, Ltd. All authors have approved the final version of the paper. The authors assert that all procedures contributing to this work comply with the ethical standards of the relevant national and institutional committees on human experimentation and with the Helsinki Declaration of 1975, as revised in 2008.

## Notes

### Competing Interest Statement

The authors have declared no competing interest.

## References

Arguello, P. A., & Gogos, J. A. (2008). A signaling pathway AKTing up in schizophrenia. J Clin Invest, 118(6), 2018–2021. doi:10.1172/JCI35931

Ashton, C. H. (2001). Pharmacology and effects of cannabis: a brief review. Br J Psychiatry, 178, 101–106.

Baron, R. M., & Kenny, D. A. (1986). The moderator-mediator variable distinction in social psychological research: conceptual, strategic, and statistical considerations. J Pers Soc Psychol, 51(6), 1173–1182. doi:10.1037//0022-3514.51.6.1173

Batalla, A., Crippa, J. A., Busatto, G. F., Guimaraes, F. S., Zuardi, A. W., Valverde, O., … Martin-Santos, R. (2014). Neuroimaging studies of acute effects of THC and CBD in humans and animals: a systematic review. Curr Pharm Des, 20(13), 2168–2185.

Beaulieu, J. M., Gainetdinov, R. R., & Caron, M. G. (2007). The Akt-GSK-3 signaling cascade in the actions of dopamine. Trends Pharmacol Sci, 28(4), 166–172. doi:10.1016/j.tips.2007.02.006

Beaulieu, J. M., Tirotta, E., Sotnikova, T. D., Masri, B., Salahpour, A., Gainetdinov, R. R., … Caron, M. G. (2007). Regulation of Akt signaling by D2 and D3 dopamine receptors in vivo. J Neurosci, 27(4), 881–885. doi:10.1523/JNEUROSCI.5074-06.2007

Bhattacharyya, S., Atakan, Z., Martin-Santos, R., Crippa, J. A., Kambeitz, J., Malhi, S., … McGuire, P. K. (2015). Impairment of inhibitory control processing related to acute psychotomimetic effects of cannabis. Eur Neuropsychopharmacol, 25(1), 26–37. doi:10.1016/j.euroneuro.2014.11.018

Bhattacharyya, S., Atakan, Z., Martin-Santos, R., Crippa, J. A., Kambeitz, J., Prata, D., … McGuire, P. K. (2012). Preliminary report of biological basis of sensitivity to the effects of cannabis on psychosis: AKT1 and DAT1 genotype modulates the effects of δ-9-tetrahydrocannabinol on midbrain and striatal function. Mol Psychiatry, 17(12), 1152–1155. doi:10.1038/mp.2011.187

Bhattacharyya, S., Crippa, J. A., Allen, P., Martin-Santos, R., Borgwardt, S., Fusar-Poli, P., … McGuire, P. K. (2012a). Induction of psychosis by Delta9-tetrahydrocannabinol reflects modulation of prefrontal and striatal function during attentional salience processing. Arch Gen Psychiatry, 69(1), 27–36. doi:10.1001/archgenpsychiatry.2011.161

Bhattacharyya, S., Crippa, J. A., Allen, P., Martin-Santos, R., Borgwardt, S., Fusar-Poli, P., … McGuire, P. K. (2012b). Induction of psychosis by Δ9-tetrahydrocannabinol reflects modulation of prefrontal and striatal function during attentional salience processing. Arch Gen Psychiatry, 69(1), 27–36. doi:10.1001/archgenpsychiatry.2011.161

Bhattacharyya, S., Crippa, J. A., Martin-Santos, R., Winton-Brown, T., & Fusar-Poli, P. (2009). Imaging the neural effects of cannabinoids: current status and future opportunities for psychopharmacology. Curr Pharm Des, 15(22), 2603–2614.

Bhattacharyya, S., Egerton, A., Kim, E., Rosso, L., Riano Barros, D., Hammers, A., . .. McGuire, P. (2017). Acute induction of anxiety in humans by delta-9-tetrahydrocannabinol related to amygdalar cannabinoid-1 (CB1) receptors. Sci Rep, 7(1), 15025. doi:10.1038/s41598-017-14203-4

Bhattacharyya, S., Falkenberg, I., Martin-Santos, R., Atakan, Z., Crippa, J. A., Giampietro, V., … McGuire, P. (2015). Cannabinoid modulation of functional connectivity within regions processing attentional salience. Neuropsychopharmacology, 40(6), 1343–1352. doi:10.1038/npp.2014.258

Bhattacharyya, S., Fusar-Poli, P., Borgwardt, S., Martin-Santos, R., Nosarti, C., O’Carroll, C., … McGuire, P. (2009). Modulation of mediotemporal and ventrostriatal function in humans by Delta9-tetrahydrocannabinol: a neural basis for the effects of Cannabis sativa on learning and psychosis. Arch Gen Psychiatry, 66(4), 442–451. doi:10.1001/archgenpsychiatry.2009.17

Bhattacharyya, S., Iyegbe, C., Atakan, Z., Martin-Santos, R., Crippa, J. A., Xu, X., . .. McGuire, P. K. (2014). Protein kinase B (AKT1) genotype mediates sensitivity to cannabis-induced impairments in psychomotor control. Psychol Med, 44(15), 3315–3328. doi:10.1017/S0033291714000920

Bhattacharyya, S., Morrison, P. D., Fusar-Poli, P., Martin-Santos, R., Borgwardt, S., Winton-Brown, T., … McGuire, P. K. (2010). Opposite effects of delta-9-tetrahydrocannabinol and cannabidiol on human brain function and psychopathology. Neuropsychopharmacology, 35(3), 764–774. doi:10.1038/npp.2009.184

Blasi, G., Napolitano, F., Ursini, G., Taurisano, P., Romano, R., Caforio, G., … Bertolino, A. (2011). DRD2/AKT1 interaction on D2 c-AMP independent signaling, attentional processing, and response to olanzapine treatment in schizophrenia. Proc Natl Acad Sci U S A, 108(3), 1158–1163. doi:10.1073/pnas.1013535108

Bloomfield, M. A., Ashok, A. H., Volkow, N. D., & Howes, O. D. (2016). The effects of Delta(9)-tetrahydrocannabinol on the dopamine system. Nature, 539(7629), 369–377. doi:10.1038/nature20153

Boggs, D. L., Nguyen, J. D., Morgenson, D., Taffe, M. A., & Ranganathan, M. (2018). Clinical and Preclinical Evidence for Functional Interactions of Cannabidiol and Delta(9)-Tetrahydrocannabinol. Neuropsychopharmacology, 43(1), 142–154. doi:10.1038/npp.2017.209

Bossong, M. G., Jager, G., Bhattacharyya, S., & Allen, P. (2014). Acute and non-acute effects of cannabis on human memory function: a critical review of neuroimaging studies. Curr Pharm Des, 20(13), 2114–2125.

Bossong, M. G., van Berckel, B. N., Boellaard, R., Zuurman, L., Schuit, R. C., Windhorst, A. D., … Kahn, R. S. (2009). Delta 9-tetrahydrocannabinol induces dopamine release in the human striatum. Neuropsychopharmacology, 34(3), 759–766. doi:10.1038/npp.2008.138

Bozzi, Y., Dunleavy, M., & Henshall, D. C. (2011). Cell signaling underlying epileptic behavior. Front Behav Neurosci, 5, 45. doi:10.3389/fnbeh.2011.00045

Brammer, M. J., Bullmore, E. T., Simmons, A., Williams, S. C., Grasby, P. M., Howard, R. J., … Rabe-Hesketh, S. (1997). Generic brain activation mapping in functional magnetic resonance imaging: a nonparametric approach. Magn Reson Imaging, 15(7), 763–770.

Bullmore, E., Long, C., Suckling, J., Fadili, J., Calvert, G., Zelaya, F., … Brammer, M. (2001). Colored noise and computational inference in neurophysiological (fMRI) time series analysis: resampling methods in time and wavelet domains. Hum Brain Mapp, 12(2), 61–78.

Bullmore, E. T., Brammer, M. J., Rabe-Hesketh, S., Curtis, V. A., Morris, R. G., Williams, S. C., … McGuire, P. K. (1999). Methods for diagnosis and treatment of stimulus-correlated motion in generic brain activation studies using fMRI. Hum Brain Mapp, 7(1), 38–48.

Bullmore, E. T., Suckling, J., Overmeyer, S., Rabe-Hesketh, S., Taylor, E., & Brammer, M. J. (1999). Global, voxel, and cluster tests, by theory and permutation, for a difference between two groups of structural MR images of the brain. IEEE Trans Med Imaging, 18(1), 32–42.

Colizzi, M., & Bhattacharyya, S. (2017). Does Cannabis Composition Matter? Differential Effects of Delta-9-tetrahydrocannabinol and Cannabidiol on Human Cognition. Curr Addict Rep, 4(2), 62–74. doi:10.1007/s40429-017-0142-2

Colizzi, M., & Bhattacharyya, S. (2018). Neurocognitive effects of cannabis: Lessons learned from human experimental studies. Prog Brain Res, 242, 179–216. doi:10.1016/bs.pbr.2018.08.010

Colizzi, M., Iyegbe, C., Powell, J., Blasi, G., Bertolino, A., Murray, R. M., & Di Forti, M. (2015). Interaction between DRD2 and AKT1 genetic variations on risk of psychosis in cannabis users: a case-control study. NPJ Schizophr, 1, 15025. doi:10.1038/npjschz.2015.25

Colizzi, M., McGuire, P., Giampietro, V., Williams, S., Brammer, M., & Bhattacharyya, S. (2018). Previous cannabis exposure modulates the acute effects of delta-9-tetrahydrocannabinol on attentional salience and fear processing. Experimental and Clinical Psychopharmacology.

Colizzi, M., McGuire, P., Pertwee, R. G., & Bhattacharyya, S. (2016). Effect of cannabis on glutamate signalling in the brain: A systematic review of human and animal evidence. Neurosci Biobehav Rev, 64, 359–381. doi:10.1016/j.neubiorev.2016.03.010

Colizzi, M., Weltens, N., McGuire, P., Lythgoe, D., Williams, S., Van Oudenhove, L., & Bhattacharyya, S. (2019). Delta-9-tetrahydrocannabinol increases striatal glutamate levels in healthy individuals: implications for psychosis. Mol Psychiatry. doi:10.1038/s41380-019-0374-8

Coolen, M. W., Statham, A. L., Gardiner-Garden, M., & Clark, S. J. (2007). Genomic profiling of CpG methylation and allelic specificity using quantitative high-throughput mass spectrometry: critical evaluation and improvements. Nucleic Acids Res, 35(18), e119. doi:10.1093/nar/gkm662

Crippa, J. A., Derenusson, G. N., Ferrari, T. B., Wichert-Ana, L., Duran, F. L., Martin-Santos, R., … Hallak, J. E. (2011). Neural basis of anxiolytic effects of cannabidiol (CBD) in generalized social anxiety disorder: a preliminary report. J Psychopharmacol, 25(1), 121–130. doi:10.1177/0269881110379283

Di Forti, M., Iyegbe, C., Sallis, H., Kolliakou, A., Falcone, M. A., Paparelli, A., … Murray, R. M. (2012). Confirmation that the AKT1 (rs2494732) genotype influences the risk of psychosis in cannabis users. Biol Psychiatry, 72(10), 811–816. doi:10.1016/j.biopsych.2012.06.020

ElSohly, M. A., Radwan, M. M., Gul, W., Chandra, S., & Galal, A. (2017). Phytochemistry of Cannabis sativa L. Prog Chem Org Nat Prod, 103, 1–36. doi:10.1007/978-3-319-45541-9_1

Emamian, E. S., Hall, D., Birnbaum, M. J., Karayiorgou, M., & Gogos, J. A. (2004). Convergent evidence for impaired AKT1-GSK3beta signaling in schizophrenia. Nat Genet, 36(2), 131–137. doi:10.1038/ng1296

Freeman, B., Smith, N., Curtis, C., Huckett, L., Mill, J., & Craig, I. W. (2003). DNA from buccal swabs recruited by mail: evaluation of storage effects on long-term stability and suitability for multiplex polymerase chain reaction genotyping. Behav Genet, 33(1), 67–72.

Fusar-Poli, P., Crippa, J. A., Bhattacharyya, S., Borgwardt, S. J., Allen, P., Martin-Santos, R., … McGuire, P. K. (2009). Distinct effects of {delta}9-tetrahydrocannabinol and cannabidiol on neural activation during emotional processing. Arch Gen Psychiatry, 66(1), 95–105. doi:10.1001/archgenpsychiatry.2008.519

Giovannetti, E., Zucali, P. A., Peters, G. J., Cortesi, F., D’Incecco, A., Smit, E. F., … Tibaldi, C. (2010). Association of polymorphisms in AKT1 and EGFR with clinical outcome and toxicity in non-small cell lung cancer patients treated with gefitinib. Mol Cancer Ther, 9(3), 581–593. doi:10.1158/1535-7163.MCT-09-0665

Hanus, L. O., Meyer, S. M., Munoz, E., Taglialatela-Scafati, O., & Appendino, G. (2016). Phytocannabinoids: a unified critical inventory. Nat Prod Rep, 33(12), 1357–1392. doi:10.1039/c6np00074f

Harris, S. L., Gil, G., Robins, H., Hu, W., Hirshfield, K., Bond, E., … Levine, A. J. (2005). Detection of functional single-nucleotide polymorphisms that affect apoptosis. Proc Natl Acad Sci U S A, 102(45), 16297–16302. doi:10.1073/pnas.0508390102

Hayasaka, S., & Nichols, T. E. (2003). Validating cluster size inference: random field and permutation methods. Neuroimage, 20(4), 2343–2356.

Hindocha, C., Quattrone, D., Freeman, T. P., Murray, R. M., Mondelli, V., Breen, G., … Di Forti, M. (2020). Do AKT1, COMT and FAAH influence reports of acute cannabis intoxication experiences in patients with first episode psychosis, controls and young adult cannabis users? Transl Psychiatry, 10(1), 143. doi:10.1038/s41398-020-0823-9

Kay, S. R., Fiszbein, A., & Opler, L. A. (1987). The positive and negative syndrome scale (PANSS) for schizophrenia. Schizophr Bull, 13(2), 261–276.

Lai, W. S., Xu, B., Westphal, K. G., Paterlini, M., Olivier, B., Pavlidis, P., … Gogos, J. A. (2006). Akt1 deficiency affects neuronal morphology and predisposes to abnormalities in prefrontal cortex functioning. Proc Natl Acad Sci U S A, 103(45), 16906–16911. doi:10.1073/pnas.0604994103

Laruelle, M., & Abi-Dargham, A. (1999). Dopamine as the wind of the psychotic fire: new evidence from brain imaging studies. J Psychopharmacol, 13(4), 358–371. doi:10.1177/026988119901300405

Mathew, R. J., Wilson, W. H., Humphreys, D. F., Lowe, J. V., & Wiethe, K. E. (1992). Regional cerebral blood flow after marijuana smoking. J Cereb Blood Flow Metab, 12(5), 750–758. doi:10.1038/jcbfm.1992.106

Matsuda, S., Ikeda, Y., Murakami, M., Nakagawa, Y., Tsuji, A., & Kitagishi, Y. (2019). Roles of PI3K/AKT/GSK3 Pathway Involved in Psychiatric Illnesses. Diseases, 7(1). doi:10.3390/diseases7010022

McLellan, A. T., Luborsky, L., Woody, G. E., & O’Brien, C. P. (1980). An improved diagnostic evaluation instrument for substance abuse patients. The Addiction Severity Index. J Nerv Ment Dis, 168(1), 26–33.

Moreira, F. A., & Wotjak, C. T. (2010). Cannabinoids and anxiety. Curr Top Behav Neurosci, 2, 429–450.

Morgan, C. J., Freeman, T. P., Powell, J., & Curran, H. V. (2016). AKT1 genotype moderates the acute psychotomimetic effects of naturalistically smoked cannabis in young cannabis smokers. Transl Psychiatry, 6, e738. doi:10.1038/tp.2015.219

Norris, H. (1971). The action of sedatives on brain stem oculomotor systems in man. Neuropharmacology, 10(21), 181–191.

Ouellet-Morin, I., Wong, C. C., Danese, A., Pariante, C. M., Papadopoulos, A. S., Mill, J., & Arseneault, L. (2013). Increased serotonin transporter gene (SERT) DNA methylation is associated with bullying victimization and blunted cortisol response to stress in childhood: a longitudinal study of discordant monozygotic twins. Psychol Med, 43(9), 1813–1823. doi:10.1017/S0033291712002784

Ozaita, A., Puighermanal, E., & Maldonado, R. (2007). Regulation of PI3K/Akt/GSK-3 pathway by cannabinoids in the brain. J Neurochem, 102(4), 1105–1114. doi:10.1111/j.1471-4159.2007.04642.x

Pertwee, R. G. (2008a). The diverse CB1 and CB2 receptor pharmacology of three plant cannabinoids: delta9-tetrahydrocannabinol, cannabidiol and delta9-tetrahydrocannabivarin. Br J Pharmacol, 153(2), 199–215. doi:10.1038/sj.bjp.0707442

Pertwee, R. G. (2008b). Ligands that target cannabinoid receptors in the brain: from THC to anandamide and beyond. Addict Biol, 13(2), 147–159. doi:10.1111/j.1369-1600.2008.00108.x

Phan, K. L., Angstadt, M., Golden, J., Onyewuenyi, I., Popovska, A., & de Wit, H. (2008). Cannabinoid modulation of amygdala reactivity to social signals of threat in humans. J Neurosci, 28(10), 2313–2319. doi:10.1523/JNEUROSCI.5603-07.2008

Qiao, X., Gai, H., Su, R., Deji, C., Cui, J., Lai, J., & Zhu, Y. (2018). PI3K-AKT-GSK3beta-CREB signaling pathway regulates anxiety-like behavior in rats following alcohol withdrawal. J Affect Disord, 235, 96–104. doi:10.1016/j.jad.2018.04.039

Sami, M. B., Rabiner, E. A., & Bhattacharyya, S. (2015). Does cannabis affect dopaminergic signaling in the human brain? A systematic review of evidence to date. Eur Neuropsychopharmacol, 25(8), 1201–1224. doi:10.1016/j.euroneuro.2015.03.011

Sánchez, M. G., Ruiz-Llorente, L., Sánchez, A. M., & Díaz-Laviada, I. (2003). Activation of phosphoinositide 3-kinase/PKB pathway by CB(1) and CB(2) cannabinoid receptors expressed in prostate PC-3 cells. Involvement in Raf-1 stimulation and NGF induction. Cell Signal, 15(9), 851–859.

Shum, C., Dutan, L., Annuario, E., Warre-Cornish, K., Taylor, S. E., Taylor, R. D., … Srivastava, D. P. (2020). Delta(9)-tetrahydrocannabinol and 2-AG decreases neurite outgrowth and differentially affects ERK1/2 and Akt signaling in hiPSC-derived cortical neurons. Molecular and Cellular Neuroscience, 103, 103463. doi:10.1016/j.mcn.2019.103463

Spielberger, C. (1983). Manual of the State-Trait Anxiety Inventory. Palo Alto, CA: Consulting Psychologists Press Inc.

Suzuki, M. M., & Bird, A. (2008). DNA methylation landscapes: provocative insights from epigenomics. Nat Rev Genet, 9(6), 465–476. doi:10.1038/nrg2341

Szutorisz, H., & Hurd, Y. L. (2016). Epigenetic Effects of Cannabis Exposure. Biol Psychiatry, 79(7), 586–594. doi:10.1016/j.biopsych.2015.09.014

Talairach, J., & Tournoux, P. (1988). [Co-planar Stereotaxic Atlas of the Human Brain.]. New York: Thieme Medical

Tan, H. Y., Nicodemus, K. K., Chen, Q., Li, Z., Brooke, J. K., Honea, R., … Weinberger, D. R. (2008). Genetic variation in AKT1 is linked to dopamine-associated prefrontal cortical structure and function in humans. J Clin Invest, 118(6), 2200–2208. doi:10.1172/JCI34725

Thirion, B., Pinel, P., Meriaux, S., Roche, A., Dehaene, S., & Poline, J. B. (2007). Analysis of a large fMRI cohort: Statistical and methodological issues for group analyses. Neuroimage, 35(1), 105–120. doi:10.1016/j.neuroimage.2006.11.054

UNODC. (2019) World Drug Report. Retrieved from https://wdr.unodc.org/wdr2019/.

van der Knaap, L. J., Schaefer, J. M., Franken, I. H., Verhulst, F. C., van Oort, F. V., & Riese, H. (2014). Catechol-O-methyltransferase gene methylation and substance use in adolescents: the TRAILS study. Genes Brain Behav, 13(7), 618–625. doi:10.1111/gbb.12147

van Winkel, R., Genetic, R., & Outcome of Psychosis, I. (2011). Family-based analysis of genetic variation underlying psychosis-inducing effects of cannabis: sibling analysis and proband follow-up. Arch Gen Psychiatry, 68(2), 148–157. doi:10.1001/archgenpsychiatry.2010.152

Wong, D. F., Wagner, H. N., Tune, L. E., Dannals, R. F., Pearlson, G. D., Links, J. M., … Gjedde, A. (1986). Positron emission tomography reveals elevated D2 dopamine receptors in drug-naive schizophrenics. Science, 234(4783), 1558–1563.

